# Effects of a combined enrichment intervention on the behavioural and physiological welfare of captive Asiatic lions (*Panthera leo persica*)

**DOI:** 10.1101/2020.08.24.265686

**Authors:** Sitendu Goswami, Shiv Kumari Patel, Riyaz Kadivar, Praveen Chandra Tyagi, Pradeep Kumar Malik, Samrat Mondol

## Abstract

The endangered Asiatic lion (*Panthera leo persica*) is currently distributed as a single wild population of 670 individuals and ∼400 captive animals globally. Although the captive lions are major hope for the species’ long-term conservation through repatriation, their welfare status and management practises need research attention. To this end, we tested the efficacy of feeding, sensory and manipulable enrichment interventions on the welfare of Asiatic lions at the conservation breeding centre of Sakkarbaug Zoological Garden, Gujarat. We adopted a holistic approach by measuring physiological and behavioural responses of 35 captive Asiatic lions, divided into control (n=16) and test (n=19) groups. The test subjects approached feeding devices first and manipulable devices for a longer duration. Manipulable devices were used homogenously with two significant time peaks, but sensory devices were used sporadically throughout the day with no discernible peak usage. The control subjects remained unchanged in all welfare parameters compared to their pre-treatment levels. However, post-enrichment behavioural assessments showed higher behaviour diversity (95% increase from the baseline period), reduced enclosure zone bias (40.25% reduction) and aberrant repetitive behaviours (80.68%) in test samples. Similarly, faecal corticosterone measures showed lower stress levels in test samples (58% decrease), confirming significant improvement in all welfare indices than control groups. These results have universal applicability to assess welfare indices of other captive species in Indian zoos. We hope that the results will encourage zoo managers and regulatory agencies to improve animal welfare practices.

## 1. Introduction

Since the early Palaeolithic depictions of the “lowenmensch” by the Aurignacians (Chauvet et al., 1996), lions have been heralded across several cultures as emblems of man’s relationship with nature (McCall, 1973). Once ubiquitous across south-western Asia between Syria and parts of eastern India (Ball, 1880; Joslin, 1984), they were hunted to extinction from large parts of their range by the late 19^th^ century (Jhala et al., 2019; MacKenzie, 2017; Storey, 1991). During the 1860s, the Asiatic lion population in Gir forests, India was struggling to survive. Timely conservation measures along with a hunting ban were instrumental to the recovery of the species. Sakkarbaug zoo, founded in 1863, played a definitive role in the conservation of Asiatic lions by treating injured and diseased individuals and maintaining a viable captive stock for future repatriation. Subsequent affirmative conservation actions including an ex-situ conservation breeding programme (Smith, 1984) led to the species revival from the edge of extinction. Today the extant population of more than 500 wild Asiatic lions(Gujarat Forest Department, 2015) inhabit fragmented habitats scattered over a human dominated landscape in Gujarat(Gogoi et al., 2020), Western India, along with approximately 400 Asiatic lions spread across zoos in several countries, with more than 60% of the captive population residing in Indian zoos and conservation breeding centres (Srivastav et al., 2018). The survival and proliferation of this species relies on repatriation to insulate the extant population from future stochastic extinction events (Jhala et al., 2019). Although the conservation breeding programme has been successful in maintaining a genetically diverse stock, the welfare status and incumbent management practices for the animals has not received adequate research focus (Pastorino et al., 2017; Goswami et al., 2020).

Conservation breeding programmes for endangered species should be designed to promote species-typical behaviours and cognitive plasticity for better post-release fitness (Rabin, 2003). The deleterious impacts of sterile captive environments manifest with loss of species-typical behaviours and increase in psychosomatic disorders in animals(Broom, 2011; Dawkins, 2004; Fraser, 1999; Rabin, 2003). Therefore, welfare-based management practices are vital for any successful conservation breeding programme (Swaisgood, 2010). The benefits of an individual-centric welfare evaluation (Joslin, 1984) complemented with targeted enrichment interventions has been successfully demonstrated for several captive animals viz., ursids (Carlstead et al., 1991; McGowan et al., 2010), felids (Powell, 1995; Suárez et al., 2017), canids (Cloutier and Packard, 2014; Leonard, 2008), equids (Bulens et al., 2013), small mammals (Clark and Melfi, 2012; Vargas and Anderson, 1999), reptiles (DeGregorio et al., 2017), and amphibians (Michaels et al., 2014). Apart from improving welfare, enrichment interventions have been shown to play an important positive role in increasing post-release fitness in several species (Brown et al., 2003; Rabin, 2003; Reading et al., 2013). Earlier study on welfare status of captive Asiatic lions reported that individual variations (personality and rearing history) are associated with differential welfare outcomes in lions housed under similar captive environments (Goswami et al., 2020), necessitating individually tailored husbandry regimen for the animals. This study focuses on identifying the areas for improvement of the current husbandry and management practices for Asiatic lions.

Feeding, sensory and manipulable enrichments have been shown to improve the welfare of captive felids (Powell, 1995; Van Metter et al., 2008). We tested the efficacy of a combined enrichment intervention on the welfare of captive Asiatic lions at Sakkarbaug Zoological Garden (SZG). We measured several behavioural (species-typical behaviour diversity, enclosure usage and aberrant repetitive behaviours) and physiological (faecal corticosterone metabolites) welfare indices as a response to enrichment interventions. This is the first controlled trial study to incorporate both behavioural and physiological tools to measure the welfare status change in response to enrichment interventions of Asiatic lions housed at a conservation breeding centre. We hope that our findings assist in the improvement of incumbent husbandry and management practices for the species.

## 2. Methods

### 2.1. Study area

We conducted this study at the Asiatic lion conservation breeding centre at SZG. The conservation breeding programme of Asiatic lions was initiated in 1958 with nine founders and has since proliferated to house more than 47 individuals housed at SZG with several breeding pairs in other participating zoos across the world. Since SZG houses both captive-born and wild-rescued individuals, it holds intrinsic value for the conservation of Asiatic lions. During the study period, the off-display conservation breeding facility housed 47 individuals in 20 large naturalistic enclosures. A schematic map of the facility is provided in Supplementary Figure 1.

### 2.2. Subjects and housing

We selected 15 enclosures housing 35 Asiatic lions for the present study. Some of the subjects were wild-rescued (N = 19, Male = 11, Female = 8), while the rest were born in captivity (N = 16, Male = 3, Female = 13). Subjects were primarily adult animals (N = 31) with a few sub-adults (N = 4) and were randomly assigned to test and control groups to ensure uniformity of treatment. The test group consisted 19 subjects (Male = 7, Female = 12) the control group consisted 16 subjects (Male = 7, Female = 9), respectively.

Asiatic lions were housed in 15 enclosures in pair (1:1 and 0:2) (N = 9) or heterosexual (1:2) (N =6) configurations (Supplementary Table 1). All subjects were housed in the same enclosure for at least a year with the same enclosure mate and were accustomed to the enclosure and management practices. In most cases subjects were housed with related individuals. All enclosures provided adequate space (above 400m^2^/animal in general with some having 1500m^2^/animal) for the animals. The naturalistic enclosures provided subjects with sufficient natural vegetation cover, simulating the habitat of Asiatic lions while protecting subjects from visitor disturbance. Enclosures were well covered with trees and shrubs, which provided subjects with a good combination of shade and sun throughout the day. Fresh drinking water was made available inside retiring cells adjoining the paddock area. Subjects were allowed free access to the paddock area and retiring cubicles throughout the day and night. However, these enclosures offered little in terms of novelty and cognitive enrichment. Our earlier study established low enclosure space utilization, low behaviour diversity, and a high incidence of aberrant repetitive behaviours prevalent in Asiatic lions housed at SZG (Goswami et al., 2020).

A group of 10 animal keepers and contractual workers carried out all husbandry work for the lions. Keepers typically reported for duty at 0700 hours and cleaned all leftover food from retiring cells during 0730-1100 hours. The leftover food items were weighed to ascertain the food consumption by each individual. The keepers would provide the daily ration of buffalo meat to the subjects between 1700-1800 hours at the feeding cubicle, where each lion was fed separately. Keepers were also tasked with behavioural monitoring of the subjects and recorded the commencement and cessation of mating events between subjects. The zoo managers and the animal keepers met regularly to ascertain the welfare need of the captive animals based on subjective evaluations. The approach to welfare assessment was preventative and based solely on incidental keeper observations hence difficult to quantify and address. We expected to observe the effect of novel enrichment devices on the behavioural and physiological welfare indices of these captive animals.

### 2.3. Study design

We hypothesized that enrichment interventions would lead to significant improvement in behavioural (Goswami et al., 2020)and physiological welfare measures for all test subjects in contrast to the controls. We divided the subjects into control (N = 16) and treatment (N = 19) groups by a blind draw of enclosures for enrichment interventions. We conducted the study in two phases: baseline and post-enrichment. During the baseline period, we measured behaviour for 14 days/subject and collected two fresh faecal samples/subject/week. This was followed by a one-week exposure to the enrichment intervention to get test subjects accustomed to the new devices. During the testing period (14 days) we recorded the behaviour of test subjects as a response to the daily enrichment intervention. Apart from the enrichment interventions, housing and husbandry conditions remained unchanged for both groups throughout the course of the study.

### 2.4. Data collection

#### 2.4.1. Behaviour observations

Two observers (viz., first author and field assistant) collected all behaviour data, which necessitated accounting for inter-observer reliability. We created a detailed ethogram of the study subjects from one month of ad-libitum behaviour sampling and compared the behaviour data recorded from the same subject by both observers to ensure consistency (Goswami et al., 2020). We commenced data collection only after inter-observer reliability levels remained consistent across three consecutive sessions (Cronbach’s alpha >0.9). We simultaneously video recorded all behaviour sessions to aid in data entry and reduce errors. We recorded behaviour during 0500-1100, 1200-1800, and 2200-0500 hours. We used instantaneous scan sampling (Altmann, 1974)at 1-minute intervals to record all behavioural states and all occurrences of behavioural events (Goswami et al., 2020). During every six-hour data collection period, we recorded four hours of behaviour data with 30 minutes rest for observers after every hour. Data for the observer-resting period was later tabulated from the video recordings. We collected 195 ± 12.3 hours of behaviour data/subject during the study period.

We measured three behavioural welfare indices: species-typical behaviour diversity (Wemelsfelder et al., 2000), spread of participation index (Plowman, 2003) and aberrant repetitive behaviours or stereotypy (Mason & Latham, 2004) to measure the welfare of Asiatic lions (Goswami et al., 2020). We compared the behavioural welfare indices of all test and control subjects across the baseline and enrichment period. Baseline information for diversity of behaviour repertoire in control and test subjects was collected using Shannon-Weiner diversity index (SWI) (Goswami et al., 2020). We measured enclosure usage patterns of subjects based on the spread of participation index (SPI) (Plowman, 2003)to understand if the enclosures provided enough complexity to meet their welfare requirement (Plowman, 2003; Ross and Shender, 2016). We divided each enclosure (N = 15) into ten zones, i.e., three primary zones (proximal, medial, and distal), which were subdivided into three secondary zones (left, middle, right), and the tenth zone was the area adjacent to the entrance to feeding cubicles (Goswami et al., 2020). We measured the proportion of aberrant repetitive behaviours (ARBs) or stereotypy as an indicator of poor welfare and compared their prevalence between control and test subjects across treatments.

#### 2.4.2. Physiological measures

We measured faecal corticosterone metabolites of all subjects to contrast the physiological impacts of the enrichment interventions on control and test subjects. Faecal corticosterone is a reliable physiological indicator of stress and compromised welfare in captive felids (Ruskell et al., 2015; Schildkraut, 2016; Vaz et al., 2017; Young et al., 2004). We collected two fresh faecal samples/week from each subject (both control and test) during the entire study period (pre-enrichment and post-enrichment).

We collected all fresh faecal samples using the dry sampling approach (Biswas et al., 2019) and stored in −20°C freezer onsite and later transported to the Wildlife Institute of India in dry ice. The samples were stored in the laboratory at −20°C freezer until further processing. To control for effects of moisture and diet we pulverized the frozen samples before lyophilizing (#FD-5, Allied Frost, New Delhi, India) them for 72 hours prior to hormone extraction (Mondol et al., 2020). Subsequently, we sieved lyophilized samples through a 0.5 mm stainless steel mesh to obtain homogenized faecal powder. We thoroughly mixed the dried faecal powder and extracted hormone by pulse-vortexing 0.1 grams of powder in 15 ml of 70% ethanol, followed by centrifugation at 2200 rpm for 20 min (Mondol et al., 2020; Wasser et al., 2010). We collected and stored the hormone extracts in 2 ml cryochill vials (1:15 dilution) and stored in −20°C freezer till further analyses.

We used corticosterone EIA kit (#K014, Arbor Assays, MI, USA) for corticosterone metabolite estimation in faecal hormone extracts. Sample extracts were air-dried inside an incubator and resuspended in assay buffer as per required dilutions. Each sample was assayed in duplicate using kit protocol and the optical density was measured at 450 nm using an ELISA plate reader (#GMB-580, Genetix Biotech Asia Pvt. Ltd., New Delhi, India). Hormone metabolite concentration is interpolated using four parametric logistic (4PL) regression function in GraphPad prism software version 5 (GraphPad Software, California, USA). Cross-reactivities of the antibody are listed in Supplementary Table 2.

We tested for parallelism and accuracy to validate the corticosterone assay. We used dilutions of pooled extracts from a random combination of male and female samples (N=20) to assess reliable quantification of corticosterone at different concentrations and find optimal dilutions for final assays (at 50% binding). We plotted the relative dose against percent bound hormones for the pools and the standards and generated best-fit curve using 4PL regression, where parallel slopes indicate similar immunoreactivity at different concentrations. For accuracy test, we spiked corticosterone standards with equal volumes of diluted faecal extract of known concentration (dilution level close to 50% bound from parallelism test) and assayed with standards. We plotted the results as regression lines using observed and expected concentrations to show that faecal contaminants were not interfering with assay accuracy at the tested dilution. We calculated inter- and intra-assay coefficients of variation using repeated measures of same-pooled extract.

#### 2.4.3. Enrichment interventions

Enrichment interventions add cognitive complexity to enclosures and provide animals the opportunity to express species-typical behaviours (Mellen and Shepherdson, 1997; Skibiel et al., 2007). We used three types of enrichment devices: manipulable (Mellen and Shepherdson, 1997; Powell, 1995), sensory (Skibiel et al., 2007), and feed (Powell, 1995; Skibiel et al., 2007) (Table 1). Manipulable enrichments included hanging lion-sam balls, burlap bags, wooden perches/ platforms, wooden planks etc., whereas the sensory enrichments included olfactory augmentations such as scent trails made from the blood of buffalo, urine of unknown conspecifics, and dung of other prey species (Sambar, *Rusa unicolor*) etc. We also installed sensory enrichment (tactile) by wrapping rough coir rope on the bark of trees inside enclosures to promote auto-grooming and scent-marking behaviours in the subjects. Nutritional enrichment has been shown to garner the highest amount of attention in captive animals and can be useful for addressing neophobia to enrichment devices in shy individuals (Powell, 1995; Resende et al., 2009; Skibiel et al., 2007). We provided a range of nutritional enrichment devices, which included frozen dressed chicken, buffalo tails that were suspended by natural fibre ropes from trees at different parts of the enclosure. While manipulable and sensory enrichment devices were quasi-permanent and required little daily care, nutritional enrichment devices needed to be replenished every day. We installed and replenished enrichment devices between 0600-0700 hours at four enclosures every day. Keepers would arrive early and confine the subjects inside retiring cells with food rewards, while the enrichment interventions were placed at designated enclosures. We furnished all enclosure zones with the same combination of manipulable, sensory and nutritional enrichment.

To calculate enrichment device preferences, we collated all scans during which Asiatic lions interacted with the enrichment devices. We also recorded the type of behaviours that were performed by the subjects while interacting with the enrichment devices viz., exploration, social play, aggression, foraging and fear (Table 2). We counted the different types of behaviours performed per enrichment device by all subjects.

**Table 1:** Description of enrichment devices used in this study at enclosures

| Sl. no | Enrichment device | Enrichment type | Description |
| --- | --- | --- | --- |
| 1 | Platform | Manipulable | A log platform made to enable subjects to access vantage points within the enclosure |
| 2 | Sam ball | Manipulable | A hard-plastic ball with no holes or perforations |
| 3 | Log branches | Manipulable | Tree logs sections with bark suspended from trees |
| 4 | Lion bite ball | Manipulable | A hard-plastic ball with a rattle inside suspended by a bungee rope with canvas sheet that acts as a bite |
| 5 | Buffalo shank | Feeding | Buffalo shank suspended from a zipline |
| 6 | Chicken carcass | Feeding | Frozen Chicken carcass suspended from a tree by natural fibers |
| 7 | Buffalo tail | Feeding | Skinned buffalo tail suspended from tree branches |
| 8 | Buffalo blood | Feeding | Frozen buffalo blood cubes, provided on a clean concrete platform |
| 9 | Star anise oil mixed with garlic | Sensory | Sprayed on tree trunks or used to create scent trails to other enrichment devices |
| 10 | Prey dung | Sensory | Sambar (*Rusa unicolor*) dung smeared on tree trunks or mixed with enclosure soil at certain zones |
| 11 | Conspecific urine | Sensory | Earth drenched in the urine conspecifics from nearby enclosures smeared on tree trunks, or mixed with enclosure substrate at certain zones |

**Table 1:** Behaviours observed that are associated with different enrichment devices

| Sl. no | Behaviour | Description | Category |
| --- | --- | --- | --- |
| 1 | Aggressive Growl | Baring teeth with short vocalizations directed at conspecifics or humans, while interacting with enrichment | Aggression |
| 2 | Charge | Run up to a conspecific at a rapid short charge as a result of being challenged while using an enrichment device | Aggression |
| 3 | Strike | Hit conspecific with one paw, while defending an enrichment device | Aggression |
| 4 | Bite | Bite conspecific, while asserting control over an enrichment device | Aggression |
| 5 | Allogrooming | Lick conspecific, while interacting with enrichment device | Social |
| 6 | Nuzzling | Rub noses with conspecifics with soft grunts, and inviting them to play with enrichment device | Social |
| 7 | Roll | Rolling in the dirt with conspecific while interacting with a enrichment device | Social |
| 8 | Rub | Rub body or neck against the enrichment device | Explore |
| 9 | Sniffing | Raise head and sniff near a sensory enrichment | Explore |
| 10 | Flehmen | Bare teeth and sniff the air near an enrichment device | Explore |
| 11 | Scent mark | Spray urine or scratch the tree next to an enrichment device | Explore |
| 12 | Digging | Dig near an enrichment device | Explore |
| 13 | Scratching | Scratch the trees where enrichment devices are installed | Explore |
| 14 | Chase | Chase away conspecifics trying to use or steal enrichment device successfully or unsuccessfully. Does not result in an overtly aggressive interaction | Chase |
| 15 | Run | Run around the enclosure with the enrichment device in mouth. May or May not be chased by conspecific | Play |
| 16 | Stalk | Stalk an enrichment device | Play |
| 17 | Play | Kick around an enrichment device | Play |
| 18 | Drag | Drag enrichment device along the enclosure | Play |
| 19 | Bite | Bite enrichment device | Play |
| 20 | Eat | Eat a food based or sensory enrichment device | Forage |
| 21 | Lick | Lick enrichment devices intently | Forage |
| 22 | Neophobia | Approach enrichment device slowly and run away | Fear |
| 23 | Fearful growl | Growling at enrichment device, ears back and cowering | Fear |
| 24 | Circling | Warily circling around the enrichment device | Fear |

**Table 1:** Variations in behavioural performances in different enrichment devices used in this study.

| **Behaviour category** | **Feed enrichment (%)** | **Manipulable enrichment (%)** | **Sensory enrichment (%)** |
| --- | --- | --- | --- |
| Aggression | 2.94 | 11.51 | 0 |
| Chase | 11.54 | 1.89 | 0 |
| Explore | 3.05 | 5.95 | 11.77 |
| Fear | 0 | 1.96 | 0 |
| Forage | 11.48 | 0 | 0 |
| Play | 0 | 35.16 | 1.21 |
| Social | 1.16 | 0.35 | 0 |
| Total | 30.20 | 56.8 | 12.99 |

### 2.5. Data analysis

We analysed all data in R v3.6.2 (R Core Team, 2020) with R studio(RStudio, 2020). We used packages “tidyverse” (Wickham et al., 2019), “psych” (Revelle, 2019)and “dplyr(Wickham et al., 2020),”ggpubr”(Kassambara, 2020), “sur”(Harel, 2020) and “ggplot2” (Wickham, 2011) to calculate statistical outputs and create graphical summaries of the findings. We conducted tests for normality and ensured that variances were similar between groups to select appropriate parametrical and non-parametrical statistical analysis. We used unpaired t-test to compare the welfare indices between test and control groups during baseline and post-enrichment interventions. We used paired t-tests, to compare the differences in welfare indices within test and control groups and measured the amount of time spent by test subjects on different types of enrichment devices. Additionally, we calculated the effect size (Cohen, 1992, p. 199)of the enrichment intervention for the test group, which is scale-independent and gives a measure of the magnitude of difference between baseline and post-treatment conditions.

## 3. Results

### 3.1. Enrichment preferences of study subjects

During 14 days of enrichment intervention, we recorded 1,09,238 scans of test subjects (n=19) using enrichment devices. The test subjects used manipulable enrichment devices for longer durations (56.8%) compared to all other types viz., sensory (12.99%) and feeding (30.20%) (Table 3). It was challenging to measure the usage levels of sensory enrichments accurately and hence these values may have been under-reported in our results. When subjects were released inside the enclosures, they approached the feed-based enrichment devices first followed by manipulable and sensory devices. This trend continued throughout the intervention period. Feed-based enrichment devices were extensively used during the first two hours of the day between 0700-0900 hours (Figure 1, Table 4a) and garnered little or no interest after they were depleted. On the other hand, manipulable and sensory enrichment device usage peaked twice daily at 0700 and 1600 hours (Figure 1, Table 4a) with continued usage throughout the day.

**Table 4:** Temporal patterns of enrichment usage and behavioural categories in the test subjects (n=35) during enrichment intervention experiments. The upper part of the table presents the enrichment use pattern, whereas the lower part of the table shows behaviour categories.

| **Enrichment use pattern at temporal scale (%)** |  |  |  |  |  |  |  |
| --- | --- | --- | --- | --- | --- | --- | --- |
| **Enrichment** | **0700-0900** | **0900-1100** | **1100-1300** | **1300-1500** | **1500-1700** | **1700-1900** | **Total** |
| **Feed** | 22.13 | 8.06 | 0 | 0 | 0 | 0 | 30.2 |
| **Manipulable** | 11.269 | 6.94 | 7.02 | 23.44 | 5.66 | 2.44 | 56.8 |
| **Sensory** | 1.9 | 2.0 | 3.18 | 0.99 | 3.81 | 1.09 | 12.99 |
| **Behaviour categories at temporal scale (%)** |  |  |  |  |  |  |  |
| **Behaviours** | **0700-0900** | **0900-1100** | **1100-1300** | **1300-1500** | **1500-1700** | **1700-1900** | **Total** |
| **Aggression** | 6.78 | 1.91 | 1.97 | 0.95 | 1.91 | 1 | 14.57 |
| **Chase** | 7.69 | 4.31 | 0.92 | 0 | 0.15 | 0.30 | 13.40 |
| **Explore** | 6.35 | 2.8 | 3.71 | 2.39 | 4.11 | 1.4 | 20.78 |
| **Fear** | 0 | 0.49 | 0.16 | 0.48 | 0.16 | 0.65 | 1.96 |
| **Forage** | 7.66 | 3.83 | 0 | 0 | 0 | 0 | 11.48 |
| **Play** | 5.95 | 3.11 | 3.39 | 20.6 | 3.11 | 0.18 | 36.29 |
| **Social** | 0.86 | 0.53 | 0.08 | 0 | 0 | 0.03 | 1.52 |

**Figure 1:**
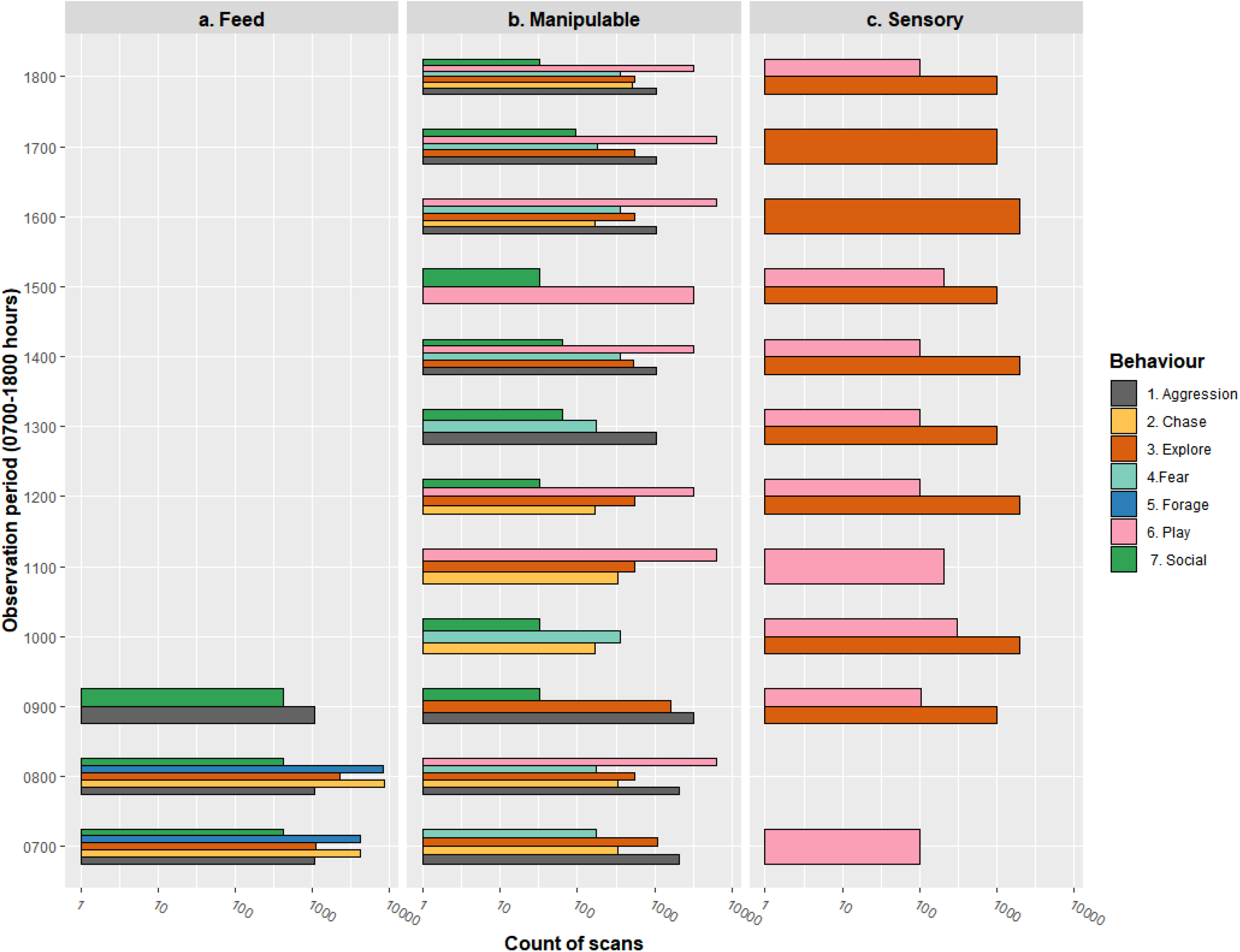
Temporal pattern of enrichment device use and behavioural patterns by Asiatic lions on (a) Feeding, (b) Manipulable, and (c) Sensory enrichment devices. Primary Y-axis represents the observation period from 0700-1800 hours. The primary x-axis represents the scan counts (log10 scale) of specific behaviours on each type of enrichment device. Hourly counts of enrichment-directed behaviours are represented as colour bars on the horizontal axis.

In terms of enrichment directed behaviours, we found that the peak of aggression (6.78%), chase (7.69%), exploratory (6.35%) and forage (7.66%) behaviours in test subjects occurred between 0700-0900 hours (Table 4b). The next peak of enrichment-directed behaviours occurred between 1300-1700 hours constituting mostly of play (20.66%) and exploratory behaviours (4.11%) (Table 4b). Early in the day (0700-0900 hours), when subjects were actively exploring around the enclosure for enrichment devices, aggressive dominant and displacement behaviours were common but subsided as the day progressed (Figure 1, Table 4b). We found that manipulable devices (11.51%) were more commonly associated with aggressive behaviours compared to food-based enrichment (2.94%) (Table 3). We also observed that positive social behaviours were more commonly associated with food-based enrichment devices (1.16%) rather than the manipulable devices (0.35%) (Table 3). Subjects usually monopolized the manipulable enrichment devices and showed high fidelity towards specific items. The feed-based enrichment devices were associated with the highest proportion of chase behaviour (11.54%), resulting in one animal monopolizing a device and chasing away others (Table 3). Some feed-based enrichment devices were consumed within minutes (e.g. dressed chicken) while others required more processing time (e.g. buffalo tail, frozen blood cubes). Once the feed enrichment devices were completely utilized, the lions concentrated on the manipulable and sensory enrichments and engaged in social and exploratory behaviours throughout the later parts of the day (Table 3, Figure 1). The manipulable enrichment devices were associated with the largest diversity of species-typical behaviours, which included 35.16% play behaviour (Table 3) and were more commonly associated with aggressive interactions (11.51%) compared to all other enrichment types. Aggressive behaviours occurred when enclosure mates tried to steal or use a certain manipulable device from the lion that was playing with it. Since we installed more enrichment devices than subjects in every enclosure, displaced individuals had the opportunity to interact with other devices when displaced by conspecifics. Sensory enrichment devices contributed primarily to exploratory behaviours (11.7%) that led to increased enclosure usage depicted by the lowering of SPI levels (Table 3).

However, it is important to point out that the true impacts of different types of enrichment devices cannot be assessed merely through the measuring the amount of time the animal spent with it, but by comparing the welfare indices of test subjects with control and their own baseline status. The results clearly demonstrate that the presence of different types of enrichment devices was instrumental in mitigating monopolization of resources by dominant individuals.

### 3.2. Behavioural welfare indices

During pre-enrichment period, behaviour diversity levels of control subjects (M = 0.9, SD = 0.3) were similar to that of the test subjects (M = 0.83, SD = 0.35). After enrichment interventions, the behaviour diversity of control subjects remained unchanged (M = 0.77, SD = 0.35, t(30)= 1.11, p = 0.29) while that of the test subjects increased significantly (M = 1.62, SD = 0.18, t(36) = −8.8, p <0.001, Cohen’s d = 2.83) (Table 5 Figure 2a).

**Table 5:** Summary statistics of welfare indices within control and test groups

| Welfare indices | Pre-enrichment | Post-enrichment | t-value* | Significance (p-value) | Effect size (Cohen's d) |
| --- | --- | --- | --- | --- | --- |
| **Control group (N=16)** |  |  |  |  |  |
| Behaviour diversity (SWI) | 0.9±0.31 | 0.77±0.35 | 1.11 | 0.29 | 0.39 |
| Spread of participation index (SPI) | 0.82±0.09 | 0.8±0.13 | 0.51 | 0.61 | 0.17 |
| Aberrant repetitive behaviours (ARB) | 12.81±6 | 13.31±6.07 | 0.23 | 0.81 | 0.08 |
| Faecal corticosterone measure | 7.1±4.08 | 6.96±3.93 | 0.09 | 0.92 | 0.03 |
| **Test group (N = 19)** |  |  |  |  |  |
| Behaviour diversity (SWI) | 0.83±0.35 | 1.62±0.18 | -8.8 | 0.0001 | 2.83 |
| Spread of participation index (SPI) | 0.77±0.12 | 0.46±0.08 | 9.37 | 0.0001 | 3.03 |
| Aberrant repetitive behaviours (ARB) | 12.32±6.03 | 2.38±2.46 | 6.65 | 0.0001 | 2.15 |
| Faecal corticosterone measure | 7.74±3.25 | 2.46±0.85 | 6.85 | 0.0001 | 2.22 |

\* For control group t(30) and test group t(36).

**Figure 2:**
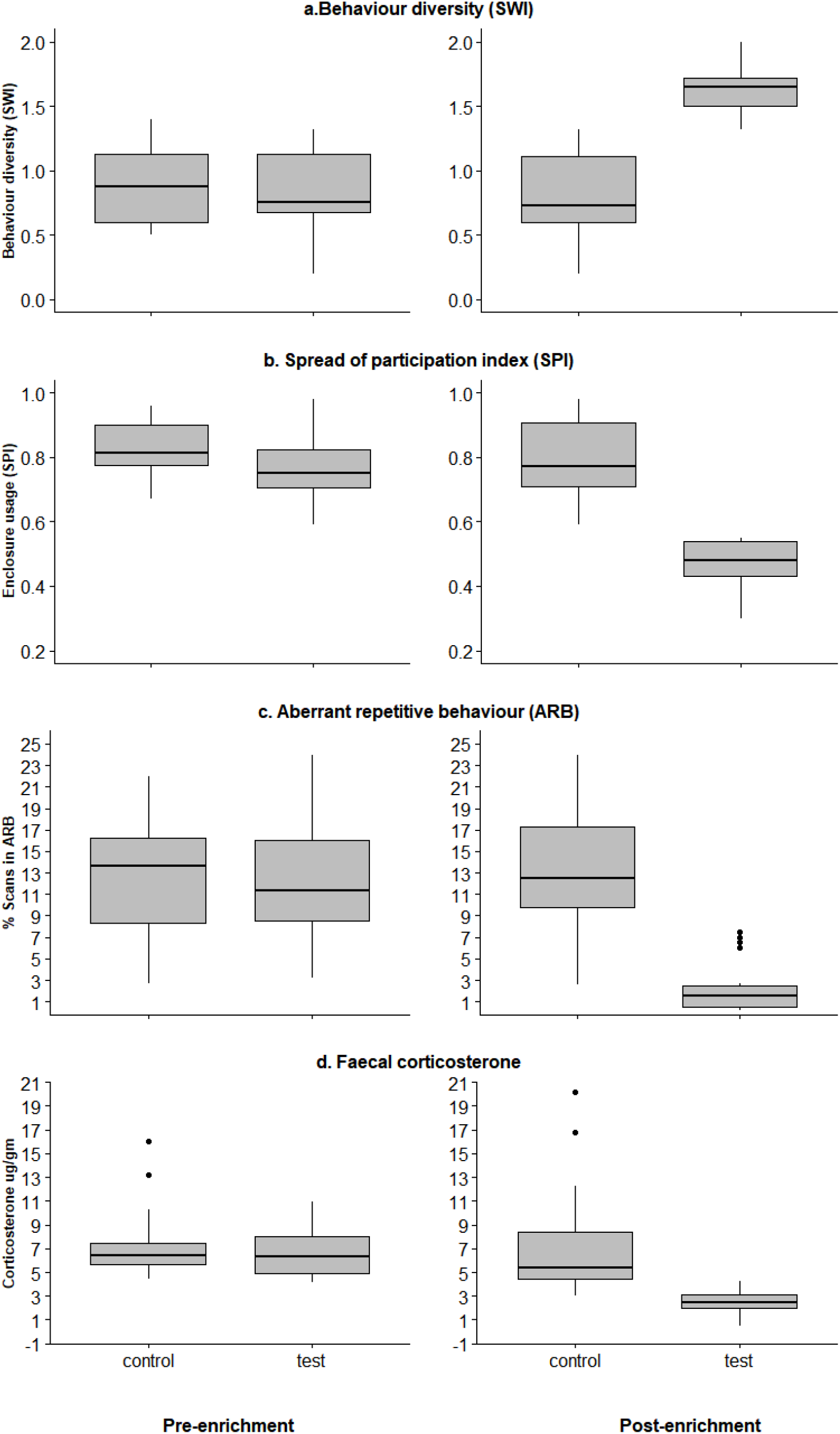
Comparison of the welfare indices between test and control groups across pre- and post-enrichment conditions. The four different welfare indices used in this study are (a) Behaviour diversity; (b) Spread of participation index; (c) Aberrant repetitive behaviours (ARB) and (d) Faecal corticosterone levels.

During the baseline, we observed high SPI values in test (M = 0.77, SD = 0.12) and control groups (M = 0.82, SD = 0.09) indicating enclosure zone use bias. However, after enrichment intervention the SPI values decreased significantly indicating, enclosure usage homogeneity (M = 0.46, SD = 0.08) for test subjects (t(36) = 9.37, p <0.0001, Cohen’s d = 3.03), while it remained unchanged for control subjects (M = 0.8, SD = 0.13, t(30)= 0.5, p = 0.61) (Table 5, Figure 2b).

During baseline, test (M = 12.32, SD = 6.03), and control subjects (M = 12.8, SD = 6) showed similar levels of aberrant repetitive behaviours. After enrichment interventions, we observed a significant decrease in ARBs in test subjects (M = 2.38, SD = 2.46, t (36) = 6.65, p < 0.0001, Cohen’s d= 2.15) while the levels remained unchanged for the control group (M = 13.31, SD = 6, t(30) = 0.23, p = 0.81) (Table 5, Figure 2c).

### 3.3. Faecal corticosterone level

Parallelism and accuracy tests for corticosterone metabolites showed reliable measures from lion faeces across different concentration ranges. Serial dilutions of faecal extracts paralleled the standard curves (Figure 3a). There were no differences between slopes of standard and pooled extract curves for corticosterone (F(1,10) = 2.06, P = 0.182), and the curves were significantly different in their elevation (F(1,11) = 66.25, P = 0.0001). Accuracy tests produced slopes of 0.92 at working dilution of 1:60 (Figure 3b), suggesting that faecal extracts did not interfere with their metabolite measurement precisions. Intra-assay coefficient of variation (CV) was 9.8%, whereas inter-assay CV was 8.99% (Supplementary Table 2).

**Figure 3:**
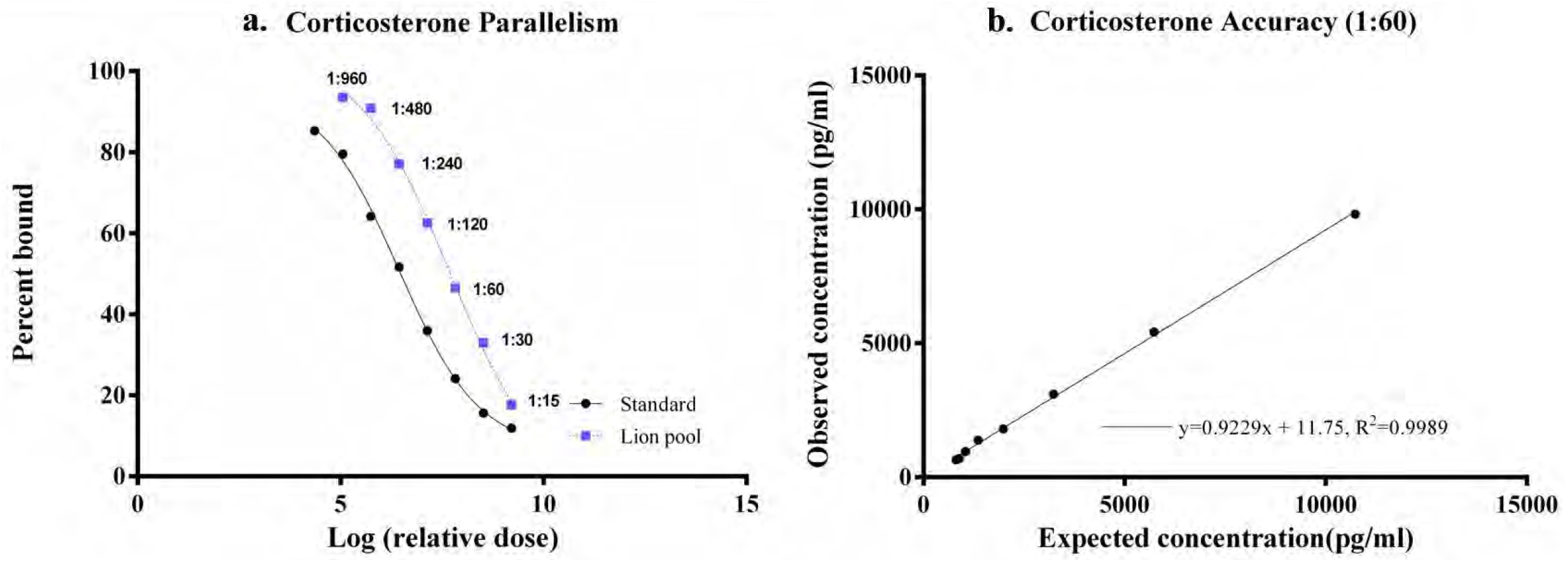
Standardization of lion faecal corticosterone through (a) parallelism and (b) accuracy tests. The graphs show accurate measures of corticosterone from captive lion faecal samples.

We compared the average faecal corticosterone levels of each subject across baseline and post-enrichment phases. Compared to baseline levels (M = 7.74 μg/gm, SD = 3.2), the faecal corticosterone measures of test subjects decreased significantly (M = 2.46 μg/gm, SD= 0.85, t(36) = 5.33, p<0.0001, Cohen’s d = 2.22), while it remained unchanged from baseline (M = 7.1 μg/gm, SD = 4.08) for control subjects (M= 6.96 μg/gm, SD = 3.93, t(30) = 0.093, p = 0.92) (Table 5, Figure 2d). These results confirm that the enrichment interventions led to significant improvement in behavioural and physiological welfare indices of test subjects as compared to the control group during pre-enrichment period.

## 4. Discussion

Animal welfare is not limited to the prevention of cruelty and amelioration of symptoms of stress, but also accords the opportunity to express species-typical behaviour patterns (Broom, 2011). The National Zoo Policy of India (MoEF,India, 1998) espouses animal welfare with captive animal management, yet welfare-centric management protocols remain divorced from husbandry practices across several zoos. Although few and far between, welfare research from Indian zoos report a prevalence of stereotypic behaviours (Goswami et al., 2020; Mallapur et al., 2007; Sanyal, 1892; Vaz et al., 2017) and elevated faecal corticosterone levels (Vaz et al., 2017) across several captive felids, and prescribe specie-appropriate enclosure design and enrichment interventions as remedial measures (Goswami et al., 2020; Mallapur et al., 2007; Vaz et al., 2017). Enrichment interventions have been shown to improve welfare conditions in several captive felids including African lions (Mellen and Shepherdson, 1997; Powell, 1995; Skibiel et al., 2007), yet enrichment protocols are seldom incorporated in the daily management of captive Asiatic lions at Indian zoos. The Asiatic lion conservation breeding programme has successfully maintained a viable captive population insulated from stochastic ecological events. Most research pertaining to captive Asiatic lions investigate their genetic and demographic status (Bagatharia et al., 2013; Pastorino et al., 2017) with only a handful mentioning their welfare (Goswami et al., 2020; Mallapur et al., 2002; Pastorino et al., 2017; Vaz et al., 2017). Our study is the first to assess the effects of enrichment protocols on the welfare of captive Asiatic lions.

Welfare encompasses both internal and external conditions affecting an animal and hence must be measured using a multifarious approach. As highlighted by Miller et al. (2020) the focus on negative welfare indicators has led to zoos adhering to minimum husbandry guidelines that attempt to suppress symptoms of poor welfare rather than trying to improve existing conditions. In this study, we measured two traditional negative welfare parameters like ARB (Mason and Latham, 2004) and faecal corticosterone levels (Schildkraut, 2016; Vaz et al., 2017) along with two positive welfare indicators (Miller et al., 2020) viz., spread of participation index (SPI) (Cabana et al., 2018; Powell, 1995) and behaviour diversity (Pastorino et al., 2017). Previous studies on captive African lions that establish the positive effects of enrichment interventions have primarily relied on behavioural welfare indices (Martínez-Macipe et al., 2015; Ncube and Ndagurwa, 2010; Powell, 1995; Regaiolli et al., 2019; Van Metter et al., 2008). Our study is the first to showcase both the behavioural and physiological impacts of enrichment interventions in captive Asiatic lions housed in a conservation breeding programme. The welfare evaluation framework used in this study can be applied universally (for other species) to measure the impacts of goal-oriented enrichment interventions.

While previous studies have tested the welfare impacts of food (Powell, 1995; Van Metter et al., 2008), manipulable (Powell, 1995), sensory (Martínez-Macipe et al., 2015; Regaiolli et al., 2019) and social stimulation (Leonard, 2008; Ncube and Ndagurwa, 2010) individually. This is the first control trial study to test the efficacy of a combined enrichment strategy on the welfare of Asiatic lions. This has allowed us to understand the temporal usage pattern of enrichment devices in Asiatic lions, which can prove useful for tailoring enrichment strategies. For example, individual test subjects showed high fidelity towards manipulable enrichment devices, which were also associated with the highest amount of conspecific-directed aggression. Therefore, to reduce stress and aggressive behaviours we recommend providing more manipulable devices than the number of animals in an enclosure. Upon release, subjects rushed to the feeding enrichment devices, therefore to prevent aggression and injury, it is important to spread such devices far apart, which will reduce chances of monopolization and food-based aggression. While sensory enrichment devices are designed to encourage exploratory behaviours in lions, the feed-based enrichment devices create a positive association for these novel objects and reduce neophobia. Finally, we found that manipulable enrichment devices allow subjects to interact and exert control over the captive environment while engaging in species-typical behaviours. Our combined enrichment strategy brought novelty in a captive environment and presented animals with the choice to express species-typical behaviour patterns. All enrichment devices used in this study can be locally sourced or fabricated with minimal effort and require less than thirty minutes to install or replenish with a team of three people.

Post-enrichment intervention, we recorded a significant improvement in the welfare indices of test subjects compared to the control subjects. Subjects that would normally stay near the retiring cells and remain inactive all day started exploring and utilizing different parts of the enclosure and followed by a significant increase in behaviour diversity levels. Faecal corticosterone levels of test subjects decreased significantly from the baseline, while it remained unchanged for control subjects, showing the inter-linkages between behavioural and physiological indices of welfare. Our findings agree with existing research that associate enrichment interventions with increased behavioural diversity (Rabin, 2003; Wemelsfelder et al., 2000), lowered SPI (Nogueira et al., 2004; Rose and Robert, 2013; Traylor-Holzer and Fritz, 1985), reduction of ARB (Clark and Melfi, 2012; Mallapur et al., 2007; Vaz et al., 2017), and reduction of stress levels (Hutchinson et al., 2012; Marcon et al., 2018; Mitra and Sapolsky, 2009; Nazar and Marin, 2011). However, it is important to point out that due to logistic constraints (sample size, time etc.) we did not address the variable responses of subjects with different personality types on enrichment interventions. Earlier work on captive Asiatic lions indicated that lions with different personality traits are likely to react differently to novel enrichment devices (Goswami et al., 2020). We suggest that future studies should focus on addressing impacts of such individual variations on differential responses and enrichment preferences with larger sample size. We studied the largest captive population of Asiatic lions, but the methodologies of this study can be scaled for smaller zoos housing fewer animals. Although we used at least three types of enrichment devices per intervention at multiple enclosure zones for experimental uniformity, at a smaller scale, zoo managers can try one enrichment device type per enclosure and rotate them with a new one every week. Our results conclusively show that enrichment interventions can lead to better behavioural and physiological welfare for Asiatic lions. We have delineated a practical but holistic approach to enrichment interventions that can be implemented and tested with little effort and manpower. With this study, we hope to empower and enable zoo managers and regulatory agencies to mandate the incorporation of enrichment interventions into daily husbandry regimen and proactively improve the welfare status of the animals under their care. We sincerely hope that the results of this study contribute to the improvement in animal welfare practices across ex-situ institutions at Indian zoos and conservation breeding programs.

## 5. Conclusion

The success of conservation breeding program depends on repatriating endangered animals to reclaimed habitats, hence housing the animals in appropriate welfare conditions while persevering behaviour diversity is of utmost importance. The Asiatic lion conservation breeding program has successfully maintained a genetically diverse stock of the species in captivity that has proliferated to an appreciable number. With the incorporation of welfare-centric management practices, the Asiatic lions housed in the program can be ideal representatives of the species and great candidates for future reintroduction programmes. Until that day of repatriation, we should strive to create a cognitively enriching environment that preserves the species-specific traits of the Asiatic lions. Like the endangered Asiatic lion, the Indian ex-situ institutions run conservation breeding programmes for several endemic species. So far, the welfare status of animals housed under conservation breeding programmes has received little research attention. We hope this study encourages zoo managers and regulators to incorporate enrichment interventions with animal management protocols.

## Ethics statement

This study was conducted in accordance with CZA norms on the welfare of animals housed in conservation breeding centres. We did not conduct any invasive sampling procedures on the subjects for experimental setup and data collection. All necessary precautions were taken to ensure that enrichment interventions were integrated with the daily husbandry regimens of the test subjects.

## Acknowledgements

We thank Dr. S.C. Pant (Principle Chief conservator of forests, Gujarat) for providing necessary permits and logistical support for this study. We acknowledge the staff members of the Sakkarbaug zoo for their help at every stage of the study. Our thanks to Mr. Salim Chuvan, head-keeper of Asiatic lions for his insights on the unique behavioural traits of the study subjects and Mr. Ashkar Bloch for his assistance during field data collection. We thank Dr. V. K. Varshney of the Chemistry and Bioprospecting Division, Forest research Institute for his laboratory facilities and Mr. Shubham Kumar for laboratory work. We also thank the Director, Dean and Research Coordinator for their continuous support. Zoological Society of London funded this study. Samrat Mondol was supported by Department of Science and Technology INSPIRE Faculty Award (IFA12-LSBM-47).

---

**Supplementary Figure 1:**
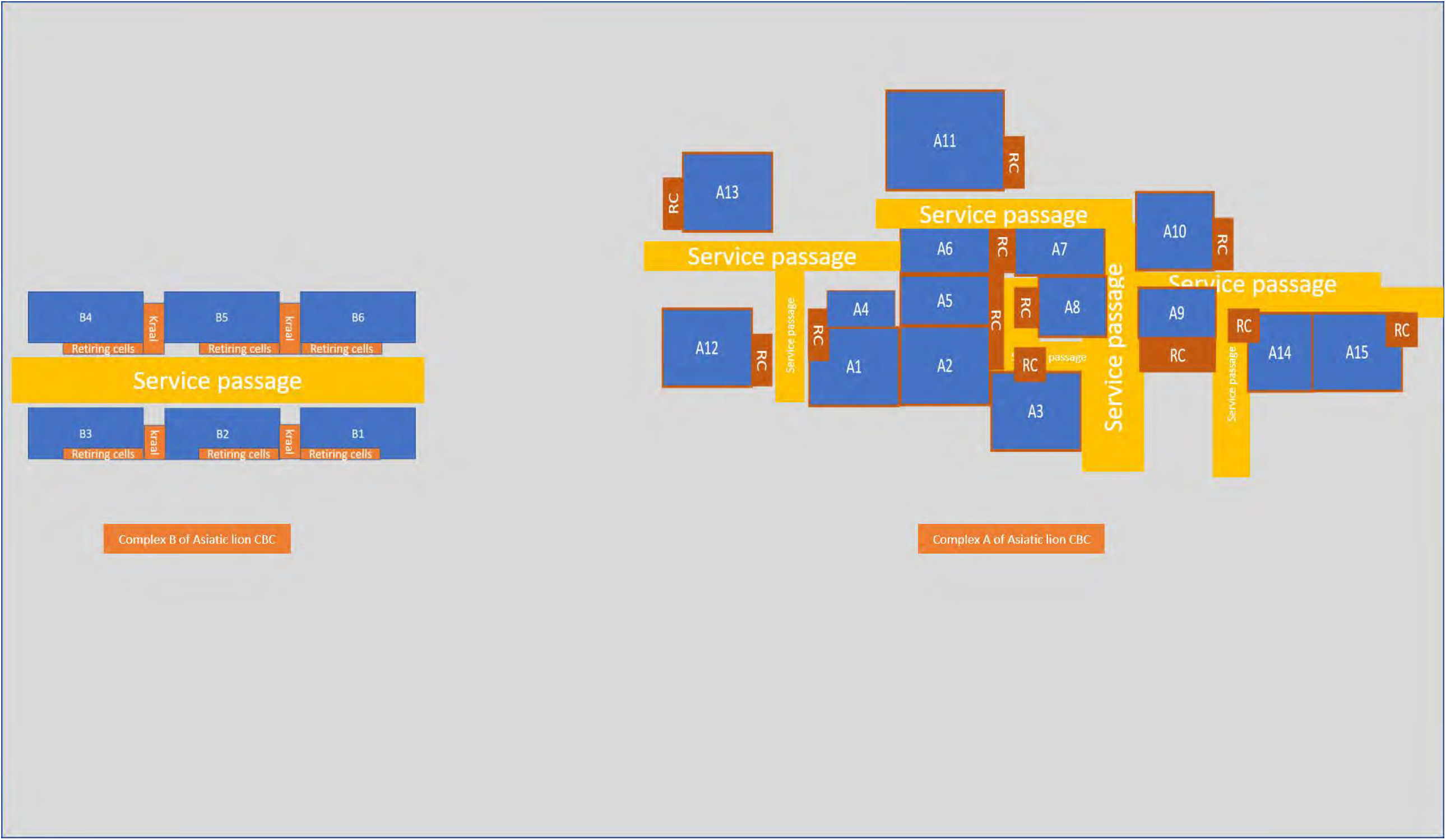
Schematic representation of enclosures at the SZG conservation breeding center for Asiatic lions.

**Table 1:** Details of subjects included in the study, viz., house name, sex (M=Male, F=Female), origin (C=Captive, W=Wild), sex ratio for housing, enclosure size, space per animal, age in days, and group.

| Sl no | Name | Sex | Origin | Sex ratio (M:F) | Enclosure size (m<sup>2</sup>) | Space/animal (m<sup>2</sup>) | Age (in days) | Group |
| --- | --- | --- | --- | --- | --- | --- | --- | --- |
| 1 | A1 | M | W | 1:1 | 1300 | 650 | 3414.00 | Control |
| 2 | Aftab | M | W | 1:1 | 1200 | 600 | 3500.00 | Control |
| 3 | Amal | M | C | 1:2 | 1576 | 788 | 2214.74 | Test |
| 4 | Ambica | F | C | 0:2 | 1123 | 561.5 | 3984.56 | Test |
| 5 | Amiya | F | C | 0:2 | 1123 | 561.5 | 2037.56 | Test |
| 6 | Ani | F | W | 1:1 | 1600 | 800 | 1946.00 | Test |
| 7 | Bahadur | M | C | 1:3 | 1600 | 533.33 | 653.33 | Test |
| 8 | AmrapurI | F | W | 1:3 | 1600 | 533.33 | 800.33 | Test |
| 9 | AmrapurII | F | W | 1:3 | 1600 | 533.3 | 800.33 | Test |
| 10 | Dhari | M | W | 1:3 | 1100 | 366.6 | 619.50 | Test |
| 11 | Dheer | M | W | 2:0 | 1600 | 800 | 5389.08 | Control |
| 12 | Gina | F | C | 1:1 | 1100 | 550 | 1316.90 | Control |
| 13 | Girm | M | W | 1:1 | 1123 | 562.5 | 3337.92 | Control |
| 14 | Hemal | M | W | 1:2 | 1700 | 566.6 | 2029.00 | Control |
| 15 | Hemali | F | W | 1:2 | 1271 | 635.5 | 2149.00 | Control |
| 16 | Jenifer | F | C | 1:1 | 1700 | 850 | 1491.33 | Test |
| 17 | Jesal | M | W | 1:2 | 1300 | 433.3 | 5722.18 | Test |
| 18 | Maheswari | F | C | 1:2 | 1300 | 433.3 | 3411.26 | Test |
| 19 | Mariyam | F | C | 1:2 | 1300 | 433.3 | 3423.35 | Test |
| 20 | Maytri | F | C | 1:1 | 1123 | 562.5 | 3072.90 | Control |
| 21 | Nagraj | M | W | 1:2 | 6542 | 2180 | 3227.00 | Control |
| 22 | Patvad | F | W | 1:1 | 1200 | 600 | 3729.00 | Control |
| 23 | Patvadm | M | W | 1:1 | 1271 | 635.5 | 1537.00 | Test |
| 24 | Radha | F | C | 1:2 | 1576 | 525.3 | 1291.30 | Test |
| 25 | Rani | F | C | 1:2 | 1576 | 525.3 | 1291.30 | Test |
| 26 | Ranita | F | C | 1:2 | 6542 | 2180 | 2119.00 | Control |
| 27 | Ranshi | F | W | 1:2 | 6542 | 2180 | 5109.00 | Control |
| 28 | Rudi | F | W | 1:1 | 1600 | 800 | 4700.00 | Control |
| 29 | Subhi | F | C | 1:1 | 1700 | 850 | 2240.33 | Test |
| 30 | Sujan | F | W | 1:1 | 1300 | 650 | 2790.80 | Control |
| 31 | Taukir | M | C | 1:1 | 1303 | 651.5 | 2448.00 | Control |
| 32 | Teeta | F | C | 1:2 | 1700 | 566.6 | 2166.93 | Test |
| 33 | Tejaswini | F | C | 1:2 | 1700 | 566.6 | 4320.93 | Test |
| 34 | Trakuda | M | W | 1:2 | 1700 | 566.6 | 3956.93 | Test |
| 35 | Veer | M | W | 2:0 | 1600 | 800 | 5387.30 | Control |

**Supplementary Table 2:** Details of the lion faecal corticosterone assay conducted in this study.

| Hormone | Assay method | Dilution | Slope (R<sup>2</sup>) | Inter-assay CV | Intra-assay CV | Cross-reactivity |
| --- | --- | --- | --- | --- | --- | --- |
| Corticosterone | EIA | 1:60 | 0.92 (0.99) | 9.0 | 9.8 | 100% with corticosterone, 12.3% with Desoxycorticosterone, 2.3% with Tetrahydrocorticosterone and <1% with Aldosterone, Cortisol, Progesterone, Dexamethasone, Corticosterone-21-Hemisuccinate, Cortisone and Estradiol |

